## Supplemental Fig 1-13 for "Deficiency in DNAH12 causes male infertility by impairing DNAH1 and DNALI1 recruitment in humans and mice"

This file includes:

**Fig S1-13**

### Fig S1

## A

| Variant filtration steps | Family1 | Family2 | Family3 |
|---|---|---|---|
| **WES sample**<br>Patients and parents or fertile brothers were sequenced. | Whole exome sequencing of the patients (P1, P2, and P3) & the mother (III:2) | Whole exome sequencing of the patients (P4, P5) & the father (III:1) & the mother (III:2) | Whole exome sequencing of the patient (P8) & the father (III:1) & the mother (III:2) & fertile brother (IV:6) |
| **Variant type**<br>Variants affect protein sequence (frameshift substitution, stop-gain, splicing, nonsynonymous SNV) were kept. | 17,549 variants | 16,898 variants | 16,837 variants |
| **Population Data (MAF)**<br>Variants with MAF>0.01 in the 1000 Genomes, ESP6500, ExAC or GnomAD were excluded, variants homozygous in our in-house 578 fertile men (41 Pakistanis, 254 Chinese and 283 Europeans) were excluded. | 1,400 variants | 1,205 variants | 1,117 variants |
| **Computational & Predictive**<br>Variants predicted to be non-deleterious by >50% software covering them were excluded. | 806 variants | 724 variants | 700 variants |
| **Expression Data**<br>Variants within genes expressed in testis were kept. | 617 variants | 514 variants | 519 variants |
| **Segregation Data**<br>Variants following inherence pattern were kept, X recessive, Autosome recessive, Compound heterozygous and Male-limited recessive models were analyzed. | XR: 0 variant;<br>AR: 2 variants;<br>Sex-limited R: 0 variant;<br>compound: 0 variant | XR: 2 variants;<br>AR: 7 variants;<br>Male-limited R: 1 variant;<br>compound: 2 variants in 1 gene | XR: 0 variant;<br>AR: 15 variants;<br>Male-limited R: 3 variants;<br>compound: 2 variants in 1 gene |
| **Functional Data**<br>Variants within genes may be functional in spermatogenesis based on SpermatogenesisOnline or literature | *DNAH12*: c. 944_945del (frameshift) | *DNAH12*: c. 164A>G (missense) | *DNAH12*: c. 2560A>G (missense) |

## B

*DNAH12* variants following AR or Sex-limited AR models are within Regions of Homozygosity (RoHs)

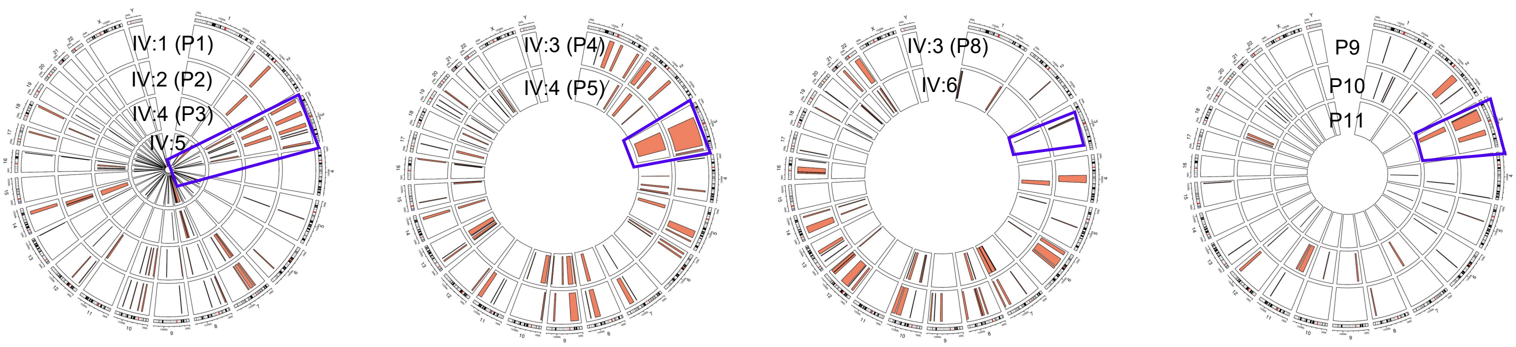

**Fig S1**

**C**

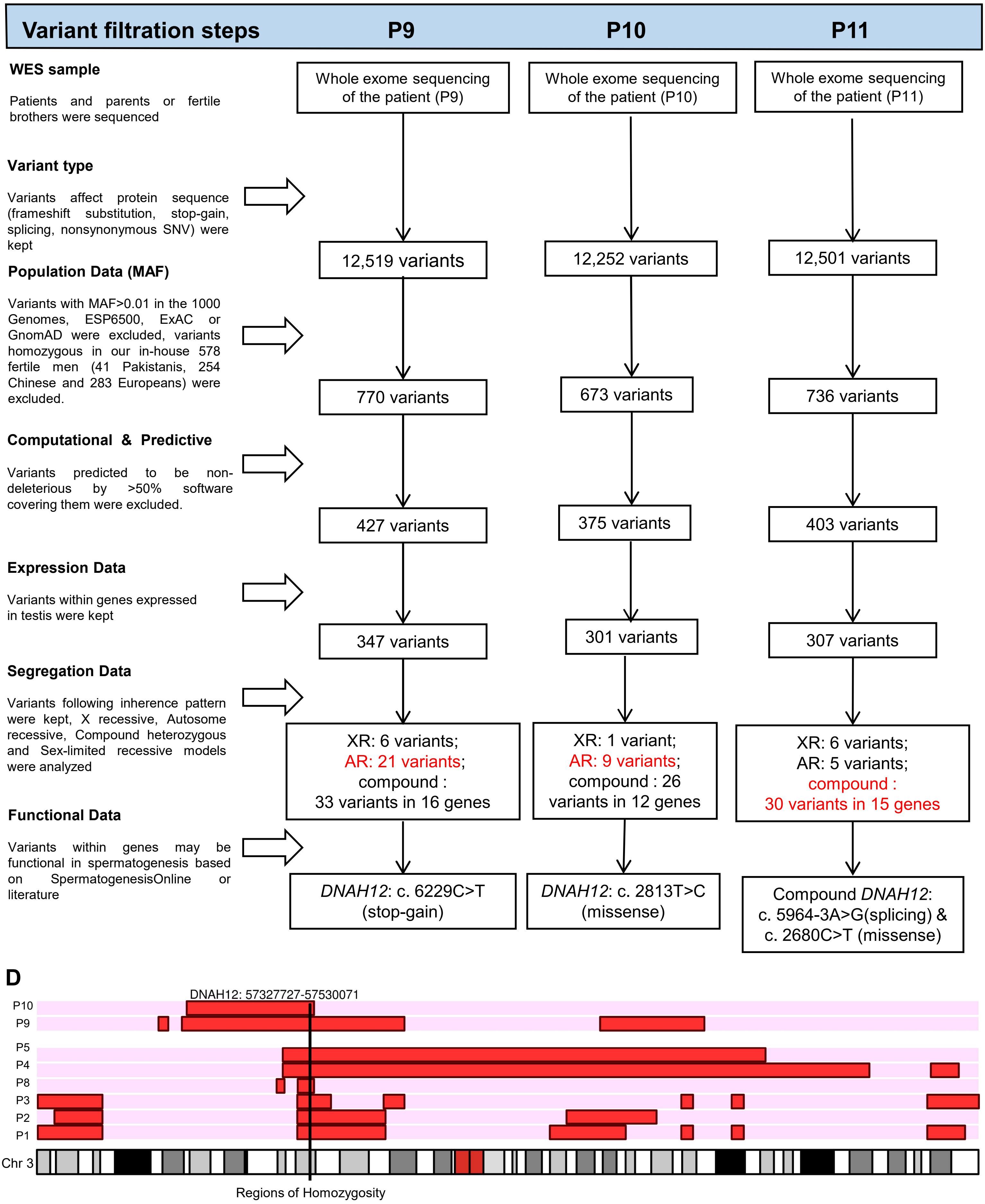

Variant filtration steps
P9
P10
P11
WES sample
Patients and parents or fertile brothers were sequenced
Whole exome sequencing of the patient (P9)
Whole exome sequencing of the patient (P10)
Whole exome sequencing of the patient (P11)
Variant type
Variants affect protein sequence (frameshift substitution, stop-gain, splicing, nonsynonymous SNV) were kept
12,519 variants
12,252 variants
12,501 variants
Population Data (MAF)
Variants with MAF>0.01 in the 1000 Genomes, ESP6500, ExAC or GnomAD were excluded, variants homozygous in our in-house 578 fertile men (41 Pakistanis, 254 Chinese and 283 Europeans) were excluded.
770 variants
673 variants
736 variants
Computational & Predictive
Variants predicted to be non-deleterious by >50% software covering them were excluded.
427 variants
375 variants
403 variants
Expression Data
Variants within genes expressed in testis were kept
347 variants
301 variants
307 variants
Segregation Data
Variants following inherence pattern were kept, X recessive, Autosome recessive, Compound heterozygous and Sex-limited recessive models were analyzed
XR: 6 variants; AR: 21 variants; compound : 33 variants in 16 genes
XR: 1 variant; AR: 9 variants; compound : 26 variants in 12 genes
XR: 6 variants; AR: 5 variants; compound : 30 variants in 15 genes
Functional Data
Variants within genes may be functional in spermatogenesis based on SpermatogenesisOnline or literature
DNAH12: c. 6229C>T (stop-gain)
DNAH12: c. 2813T>C (missense)
Compound DNAH12: c. 5964-3A>G(splicing) & c. 2680C>T (missense)
D
DNAH12: 57327727-57530071
P10
P9
P5
P4
P8
P3
P2
P1
Chr 3
Regions of Homozygosity

**Fig S1. Flowchart for analyses of the whole-exome sequencing data.** (A) and (C) *In-silico* analyses and filtering of variants in patients from Pakistani (A) and Chinese (C) families. (B) and (D) *DNAH12* variants are located in the regions of homozygosity (RoHs) of affected Pakistani or Chinese patients. RoHs are marked in red. The purple frames mark the location of the mutations.

#### Fig S2

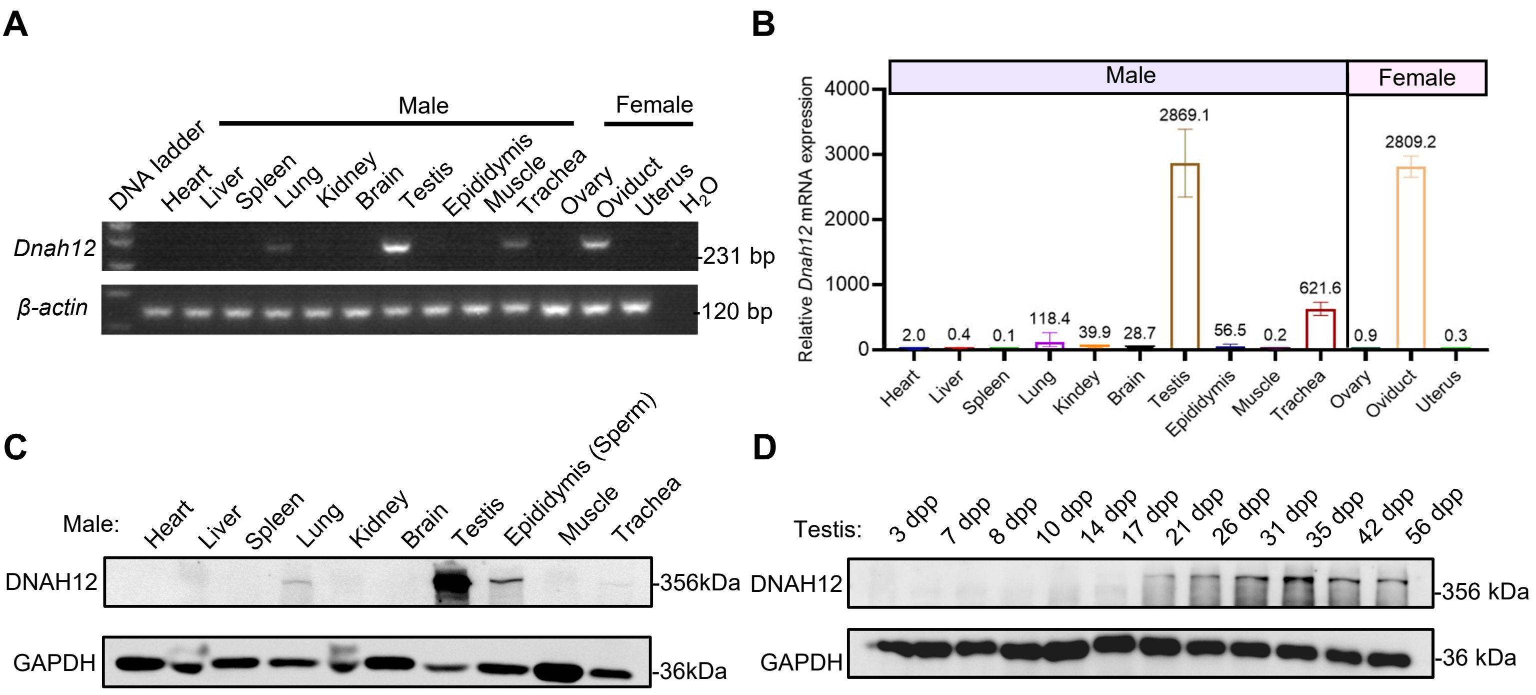


**Fig S2. Expression pattern of *Dnah12* in different mouse tissues and developmental stages of the testis.** (A) RT–PCR analysis of *Dnah12* expression in different tissues from adult mice. *β-actin* served as the reference gene. (B) Real-time quantitative PCR for *Dnah12* transcripts in various 8-week-old mouse tissues (most organs were obtained from male mice, except ovary, oviduct, and uterus which were gained from female mice), *β-actin* gene served as reference, and the experiments were repeated 3 times. (C) Immunoblotting assay of the DNAH12 expression among 10 different tissues from adult mice. (D) Immunoblotting analysis of the testicular DNAH12 expression at different days postpartum (dpp). GAPDH served as the loading control.

#### Fig S3

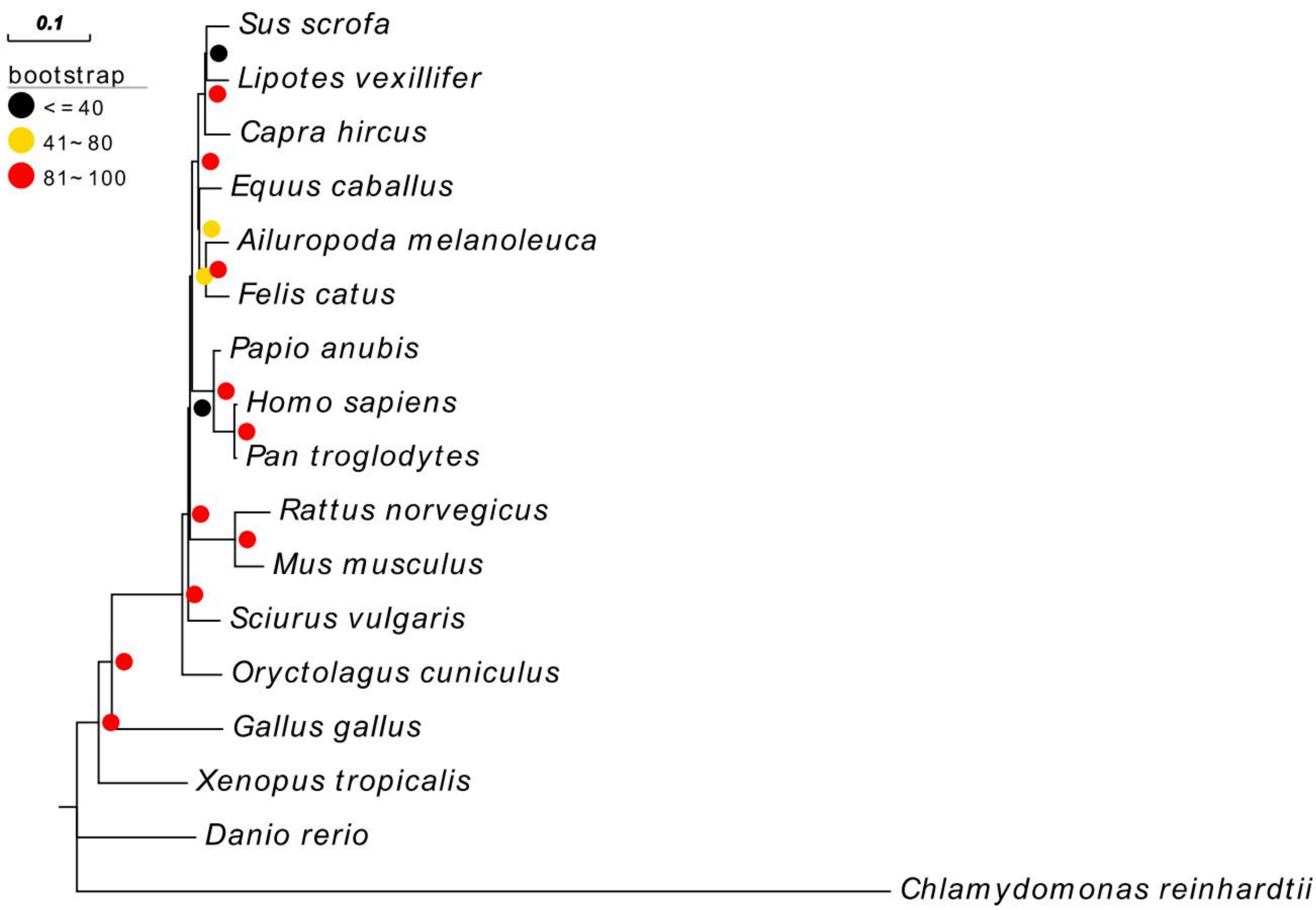


**Fig S3. Phylogenetic analysis of the DNAH12 homologous proteins in different species.**

#### Fig S4

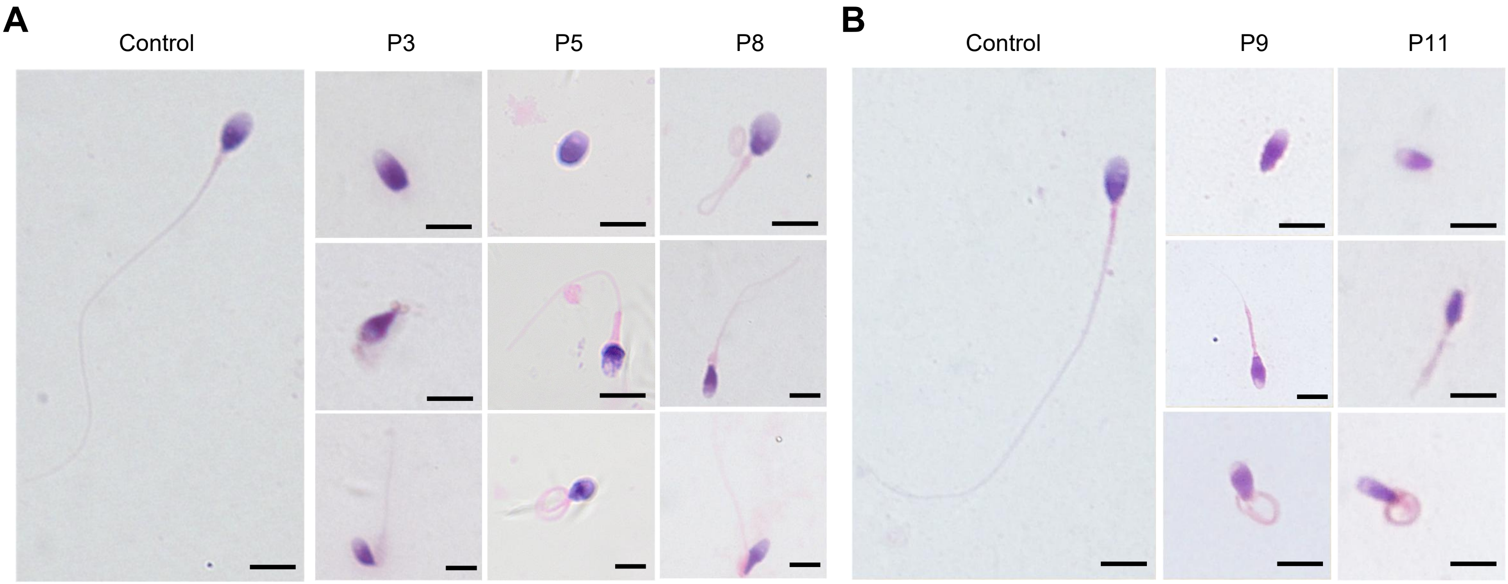


**Fig S4. Sperm morphology of fertile controls and patients.** (A) Representative micrographs from men harboring bi-allelic *DNAH12* variants (P3 of family 1, P5 of family 2, and P8 of family 3) and a fertile control by H&E staining. (B) Representative micrographs from P9, P11, and a fertile control by Papanicolaou staining. Scale bars,10 μm.

#### Fig S5

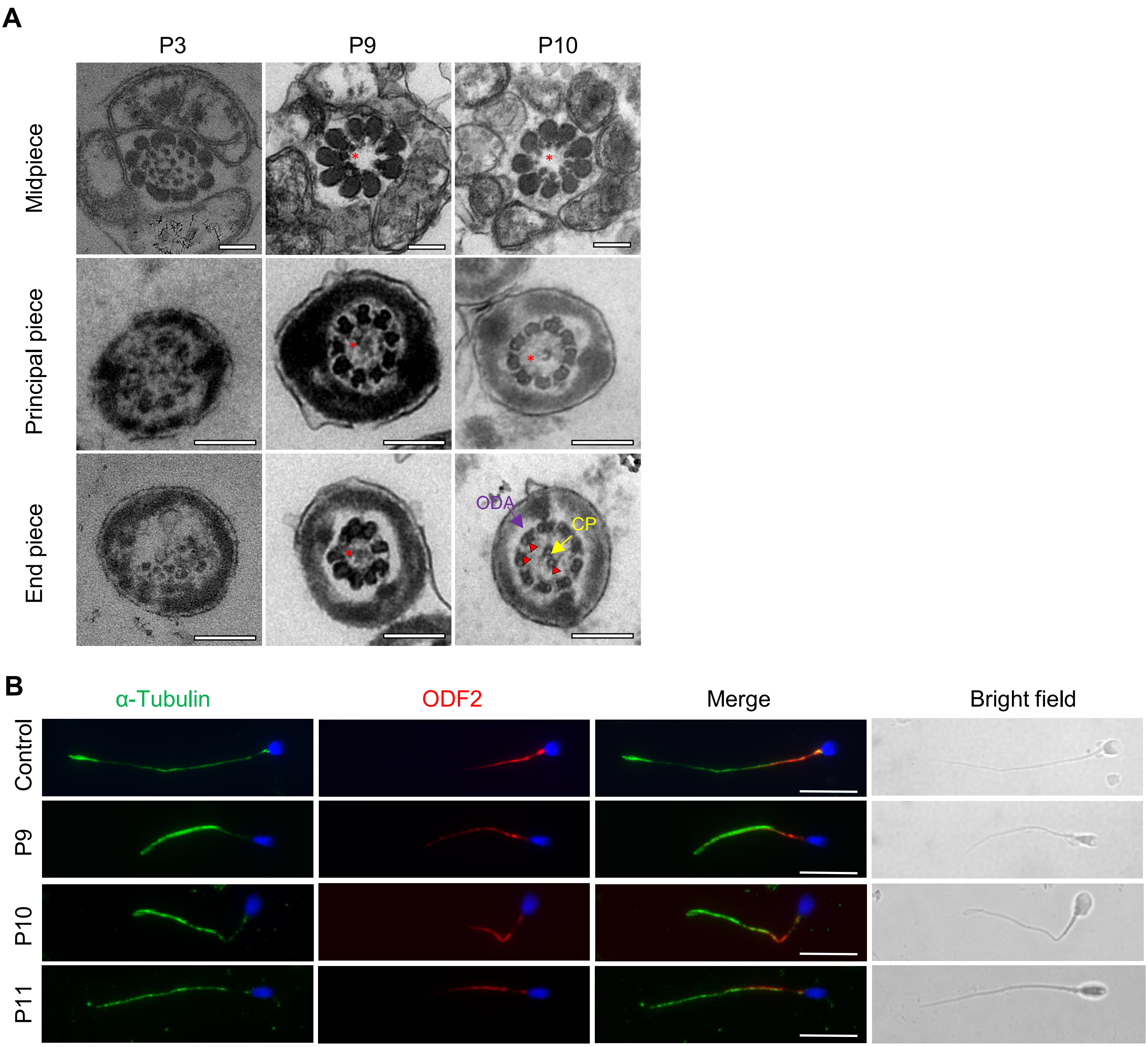


**Fig S5. TEM images of men harboring bi-allelic *DNAH12* variants and immuno-fluorescence assay of ODF2 in sperm of control and P9.** (A) Representative TEM micrographs from men harboring bi-allelic *DNAH12* variants (P3 of Famliy1, P9, P10). Red asterisks mark CP loss and red triangle marks the positions where IDAs are impaired or absent while adjacent ODAs are identifiable. Scale bars, 200 nm. (B) Representative image of spermatozoa from fertile controls and patients carrying *bi-allelic DNAH12* variants stained with ODF2 and α-Tubulin antibodies, and Hoechst 33342 (blue). Scale bars,10 μm.

#### Fig S6

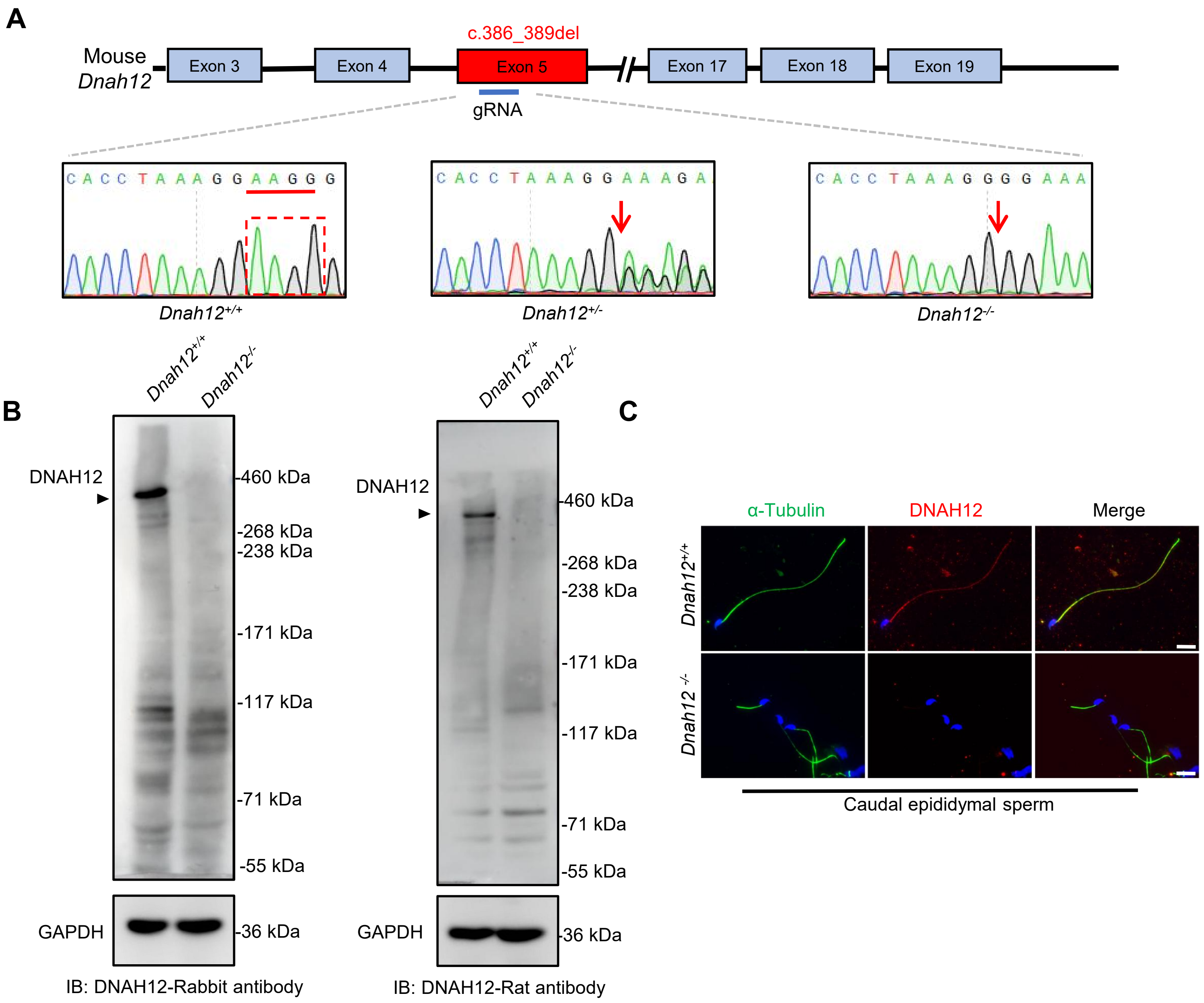


**Fig S6. Generation of *Dnah12*$^{-/-}$ mice model and validation of rabbit and rat host DNAH12 antibodies.** (A) Schematic illustrating construction of the *Dnah12*$^{-/-}$ mouse model and Sanger sequencing validation of *Dnah12*$^{+/+}$, *Dnah12*$^{+/-}$, *and Dnah12*$^{-/-}$ mice. Red arrows indicate the position where 4bp deletion occurs. (B) Immunoblotting assay of testis lysate from 10 weeks *Dnah12*$^{+/+}$ and *Dnah12*$^{-/-}$ mice using DNAH12-Rabbit antibody or DNAH12-Rat antibody. (C) Representative images of caudal epididymal sperm from *Dnah12*$^{+/+}$ and *Dnah12*$^{-/-}$ mice co-stained α-Tubulin and DNAH12 antibodies. Scale bars, 10 μm.

#### Fig S7

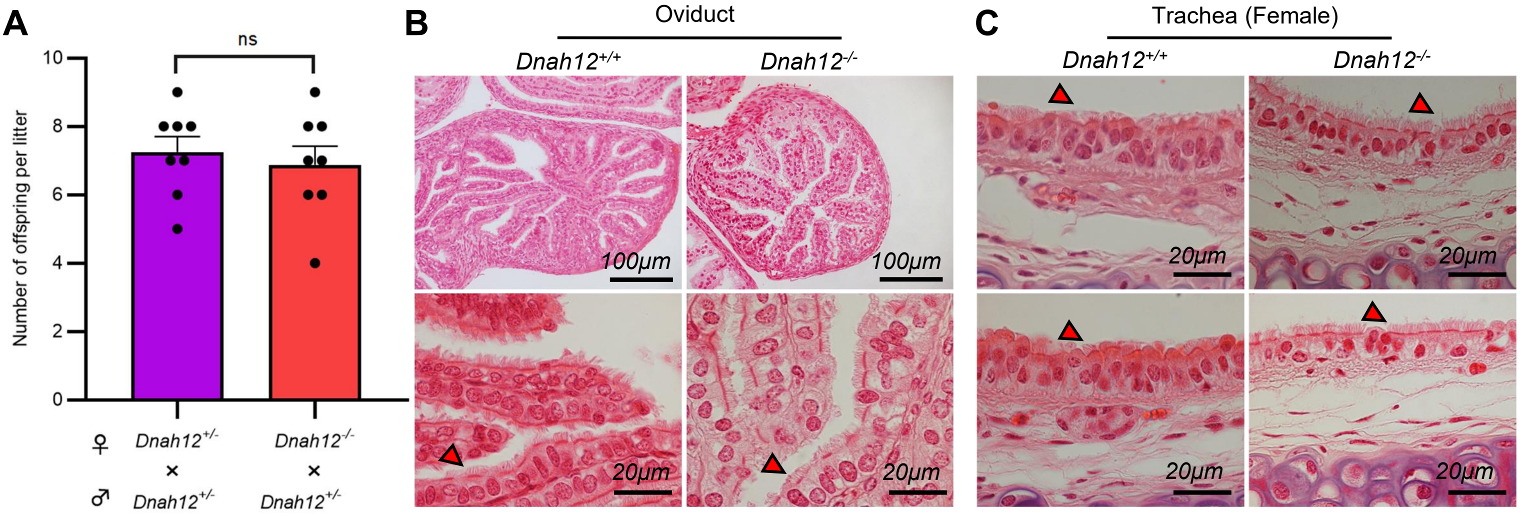


**Fig S7.Normal development and fertility in *Dnah12*$^{-/-}$ female mice.** (A) Number of offspring per litter of *Dnah12*$^{+/-}$ and *Dnah12*$^{-/-}$ *from* 8-10 weeks female mice, the two groups were caged with *Dnah12*$^{+/-}$ male mice, and the number of offspring was recorded respectively. (B-C) Oviductal (B) or tracheal (C) histology of *Dnah12*$^{+/+}$. The red triangles mark the cilia with normal morphology in *Dnah12*$^{-/-}$ 8-week-old female mice and the cilia shapes and lengths were similar in the two groups. Scare bars are shown in the figures.

### Fig S8

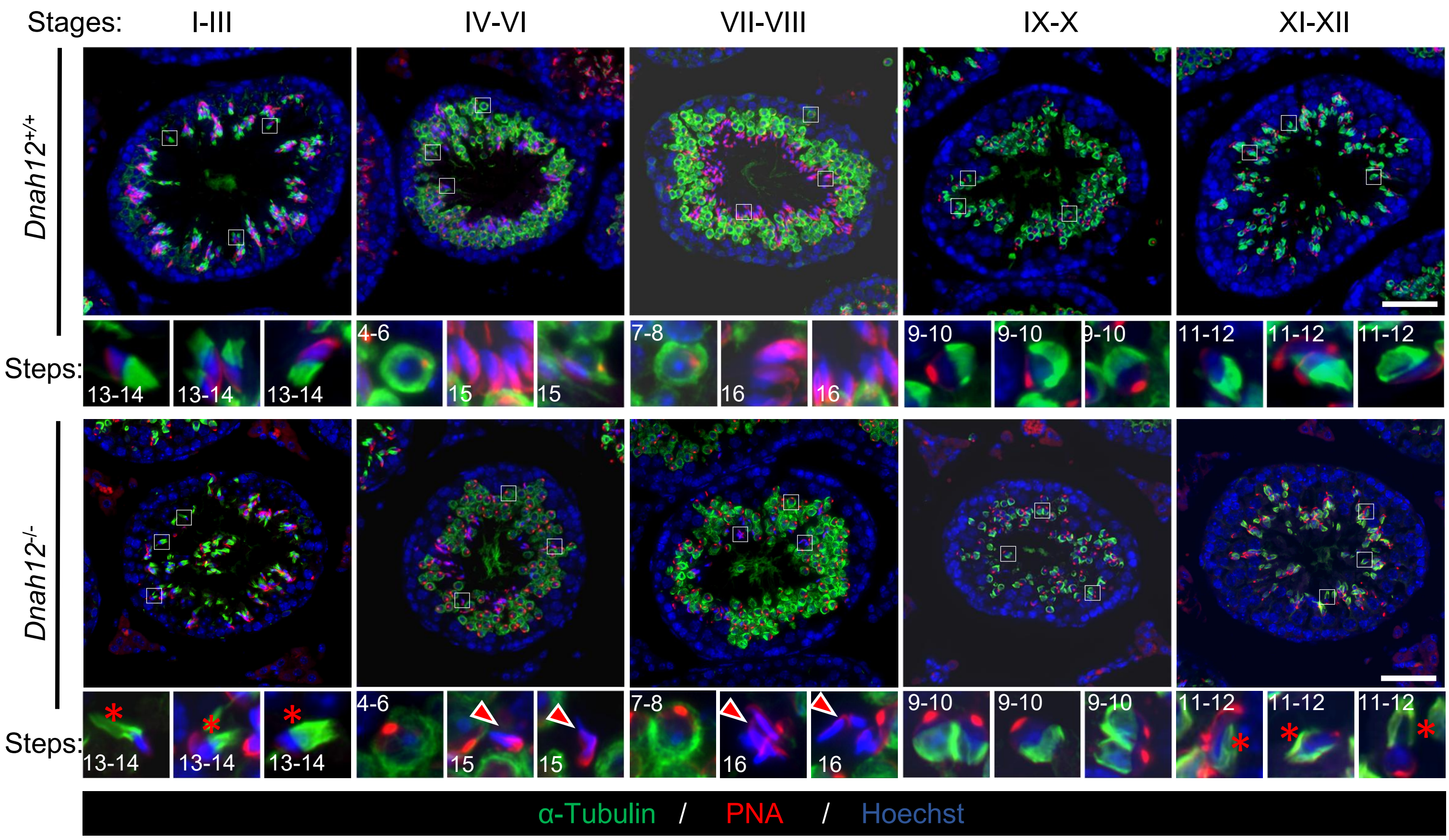


**Fig S8. Deficiency in manchette organization and abnormal sperm head shaping during spermiogenesis in *Dnah12*$^{-/-}$ mice.** Representative images of seminiferous tubule sections from 8-week-old *Dnah12*$^{+/+}$ and *Dnah12*$^{-/-}$ mice, co-stained by PNA (red) and α-Tubulin (green) antibodies. DNA was stained with Hoechst 33342. Scale bars, 50 μm.

### Fig S9

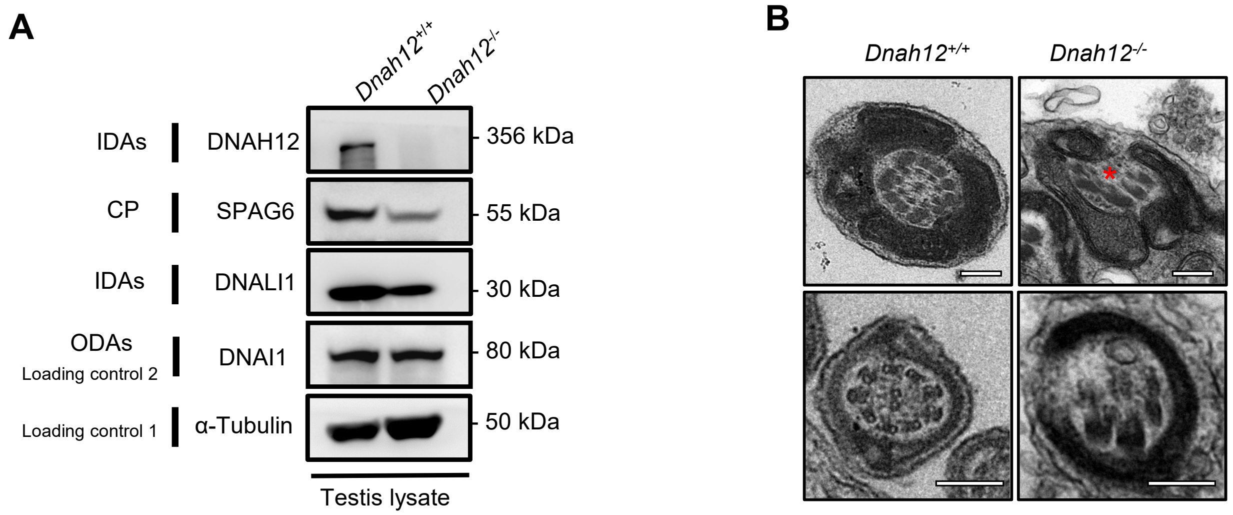


**Fig S9. The CP and DMTs structures were impaired in the *Dnah12*$^{-/-}$ testicular sperm axoneme.** (A) Immunoblotting of testis lysate from *Dnah12*$^{+/+}$ and *Dnah12*$^{-/-}$ mice using SPAG6, DNALI1 antibodies. DNAI1 and α-Tubulin were used as the loading controls. (B) Representative TEM micrographs showing cross-sections of testicular sperm axoneme from *Dnah12*$^{+/+}$ and *Dnah12*$^{-/-}$ mice. The CP and DMTs were missing or disarranged in the *Dnah12*$^{-/-}$ group. Red asterisks mark the CP loss. Scale bars, 200 nm.

#### Fig S10

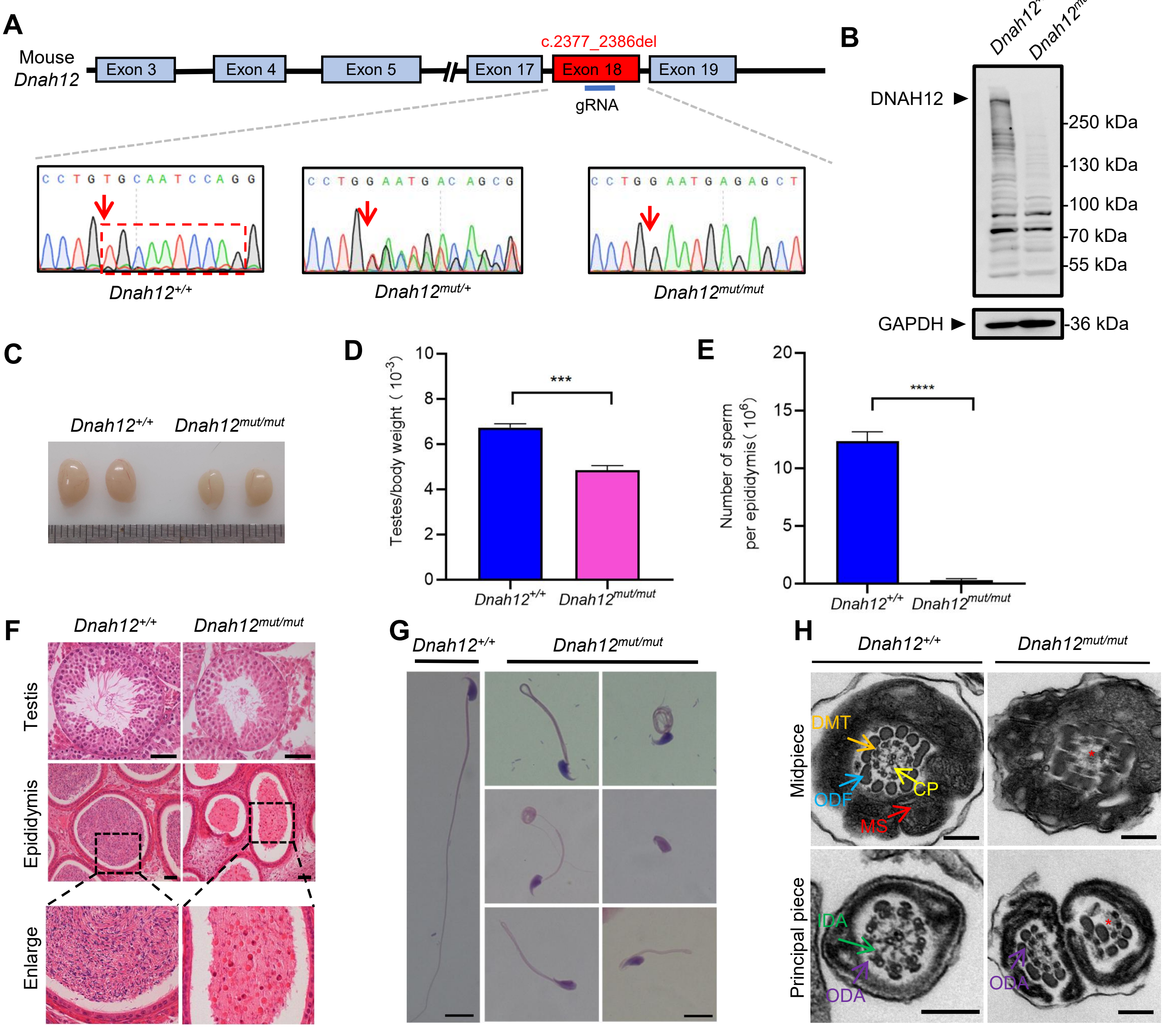

A
Mouse Dnah12
Exon 3
Exon 4
Exon 5
Exon 17
Exon 18
Exon 19
c.2377_2386del
gRNA
Dnah12+/+
Dnah12mut/+
Dnah12mut/mut
B
Dnah12+/+
Dnah12mut/mut
DNAH12
250 kDa
130 kDa
100 kDa
70 kDa
55 kDa
GAPDH
36 kDa
C
Dnah12+/+
Dnah12mut/mut
D
Testes/body weight（10-3）
***
Dnah12+/+
Dnah12mut/mut
E
Number of sperm per epididymis（106）
****
Dnah12+/+
Dnah12mut/mut
F
Dnah12+/+
Dnah12mut/mut
Testis
Epididymis
Enlarge
G
Dnah12+/+
Dnah12mut/mut
H
Dnah12+/+
Dnah12mut/mut
Midpiece
Principal piece
DMT
ODF
CP
MS
IDA
ODA

**Fig S10 Generation of *Dnah12*$^{mut/mut}$ mice.** (A) Schematic illustrating construction of the *Dnah12*$^{mut/mut}$ mice model and Sanger sequencing results of *Dnah12*$^{+/+}$, *Dnah12*$^{+/mut}$, and *Dnah12*$^{mut/mut}$ mice. The red arrows indicate the position where the mutation occurs. (B) Immunoblotting of testicular lysate from *Dnah12*$^{+/+}$ and *Dnah12*$^{mut/mut}$ mice using homemade DNAH12-Rabbit antibody. GAPDH was used as a loading control. The arrowheads indicate the target bands. (C) Representative images of testes from *Dnah12*$^{+/+}$ and *Dnah12*$^{mut/mut}$ mice. (D) Testes to body weight ratios of *Dnah12*$^{+/+}$ or *Dnah12*$^{mut/mut}$ mice. (E) Number of sperm per epididymis of *Dnah12*$^{+/+}$ and *Dnah12*$^{mut/mut}$ mice. The data were obtained from three mice for each genotype. (F) Histological sections of testis and epididymis from *Dnah12*$^{+/+}$ and *Dnah12*$^{mut/mut}$ after H&E staining. Scale bars, 50 μm. (G) Morphology of the spermatozoa obtained from *Dnah12*$^{+/+}$ mice or *Dnah12*$^{mut/mut}$ mice cauda epididymis after H&E staining. Scale bars,10 μm. (H) Representative TEM micrographs showing cross sections of sperm flagella from *Dnah12*$^{+/+}$ and *Dnah12*$^{mut/mut}$ mice. Scale bars, 200 nm.

#### Fig S11

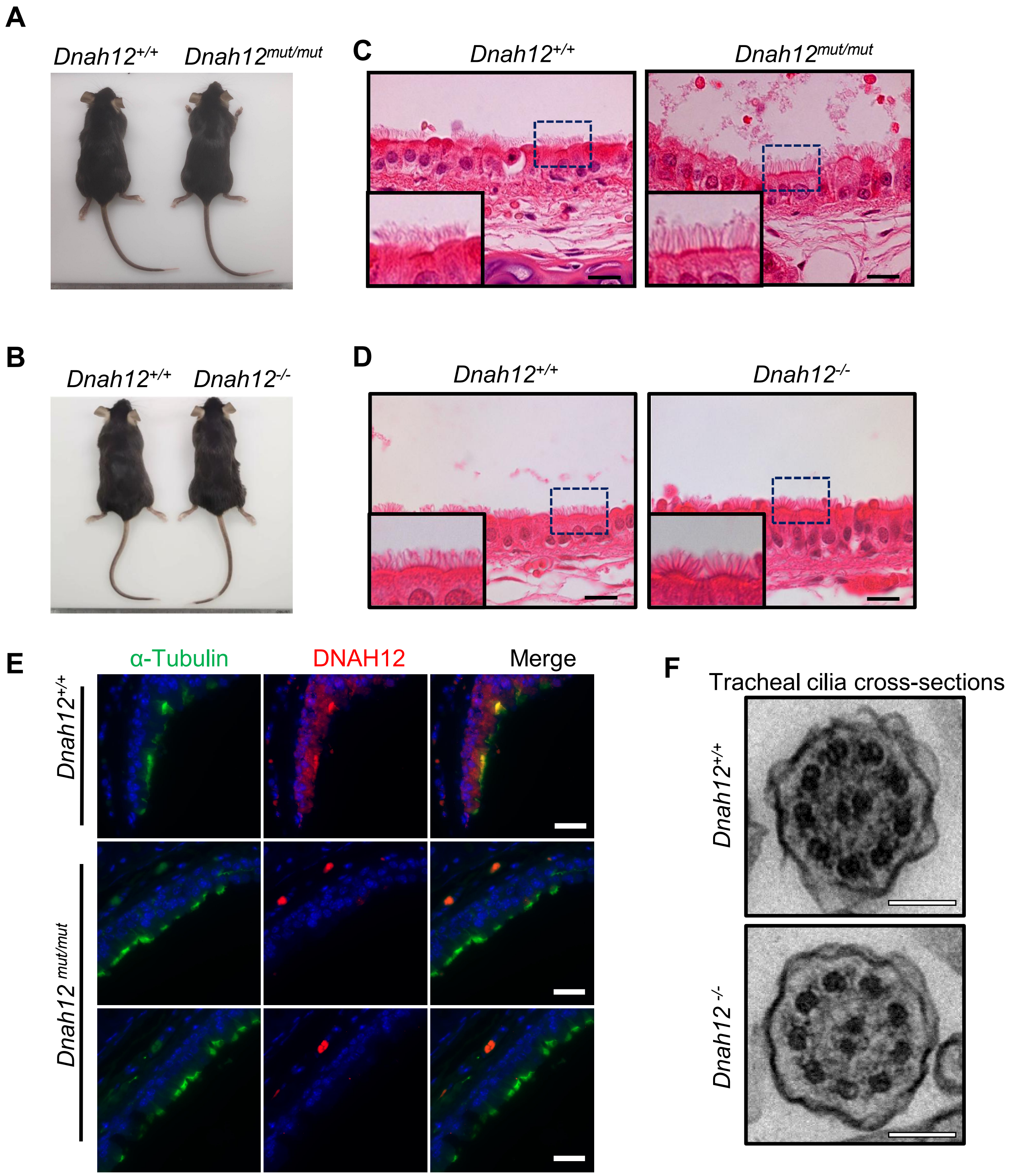


**Fig S11. Normal development and no obvious PCD symptoms were observed in *Dnah12$^{mut/mut}$* and *Dnah12$^{-/-}$* males.** (A-B) No obvious developmental abnormalities were observed in adult *Dnah12$^{mut/mut}$* (A) or *Dnah12$^{-/-}$* (B) mice. (C-D) Tracheal histology showed the cilia shape and lengths of *Dnah12$^{mut/mut}$* (C) or *Dnah12$^{-/-}$* (D) were similar to *Dnah12$^{+/+}$* in the two groups. Scare bars, 20 μm. (E) Representative images of tracheal cilia co-stained α-Tubulin (green) and DNAH12 (red) antibodies. Scale bars, 20 μm. (F) TEM micrographs showing cross sections of tracheal cilia cross-sections from *Dnah12$^{+/+}$* and *Dnah12$^{-/-}$* mice. Scale bars, 200 nm.

**Fig S12**

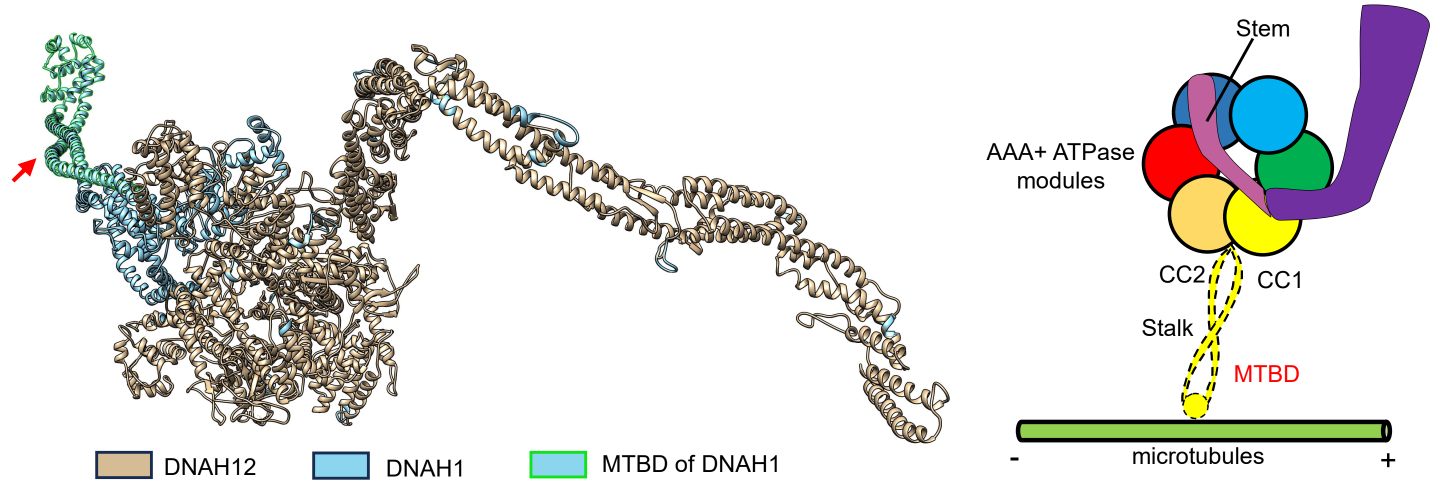


**Fig S12. DNAH12 lacks the MTBD domain.** The three-dimensional protein models of human DNAH12, DNAH1, and DNAH12 were colored golden while DNAH1 was colored celeste, for better visualization, the MTBD of DNAH1 was box selected with green lines. The proposed cartoon structures of the DNAH family are shown on the right, while DNAH12 lacks the MTBD domain.

### Figure S13

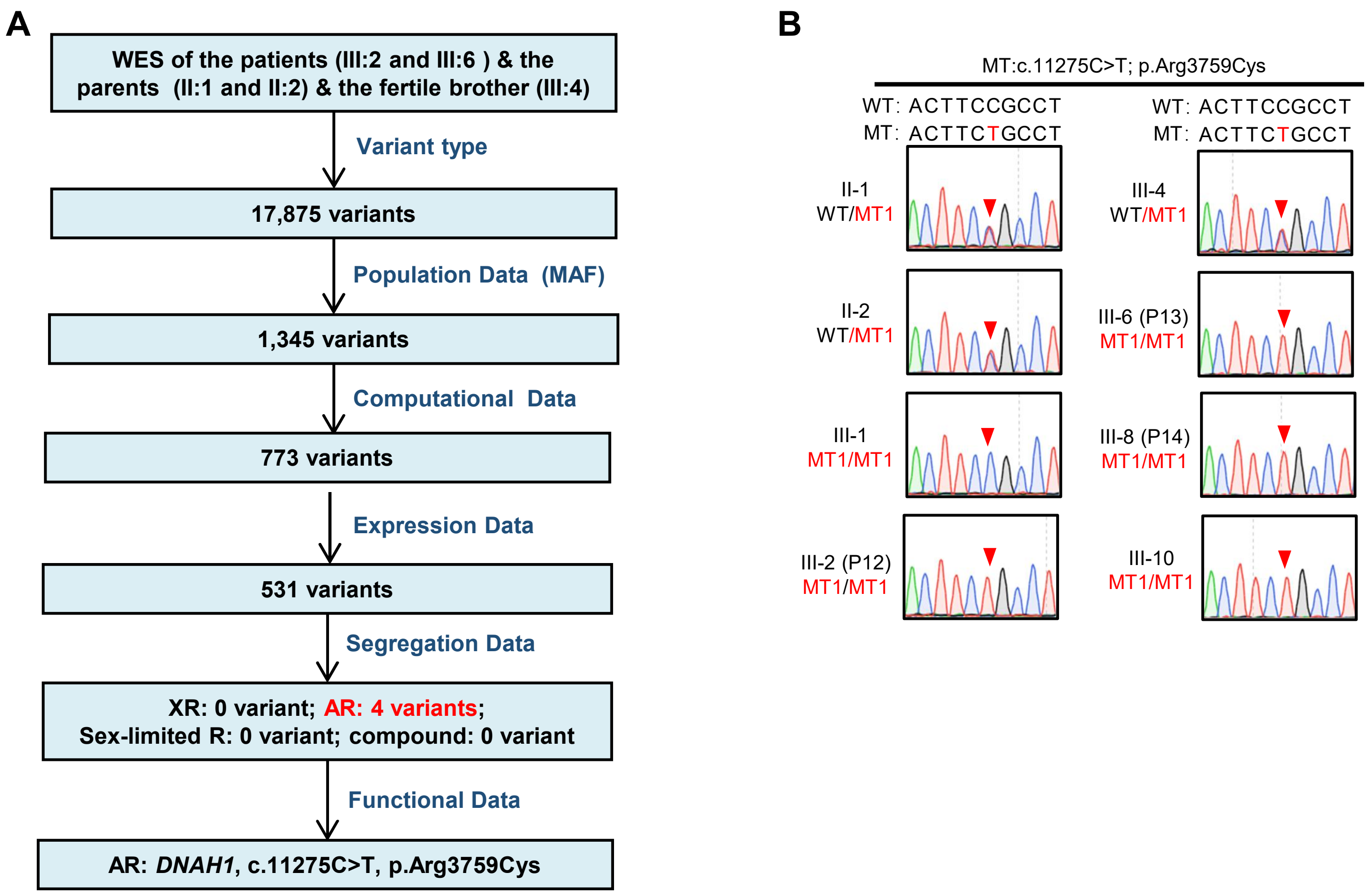


**Fig S13. Flowchart for analyses of the whole-exome sequencing of a Pakistani family and validation of *DNAH1* mutation through Sanger sequencing.** (A) Flowchart for analyses of the whole-exome sequencing of a Pakistani family with 3 infertile male patients. (B) Validation of the candidate homozygous *DNAH1 mutation* c.11275C>T, p.Arg3759Cys in the family members by Sanger sequencing.
