## Supplemental Tables 1-5 for "Deficiency in DNAH12 causes male infertility by impairing DNAH1 and DNALI1 recruitment in humans and mice"

| **Table S1. Primers used in the study** | |
| --- | --- |
| Primer names | Primer sequences (5'-3') |
| Sanger Sequencing: |  |
| Sanger-mut1-F | GTACTCTCTAAAGTAACAGTG |
| Sanger-mut1-R | GAGCTTCGGCTATTCGTTCC |
| Sanger-mut2-F | TGTCCATCTCCCAGAAAACA |
| Sanger-mut2-R | CATGAACAACCTCCACAGTT |
| Sanger-mut3-F | TTTCTTCACCATCCCGTTCT |
| Sanger-mut3-F | GAAGAAAGAATACAGACTCC |
| Sanger-mut4-F | CAAGCCCATATAATCTGACAG |
| Sanger-mut4-F | GCACAAACAGACGGATCATA |
| Sanger-mut5&mut6-F | ATACATCCCTACAGTCTCC |
| Sanger-mut5&mut6-R | AAGTCTGTATCGTGATACTG |
| Sanger-mut7-F | CTCTCCTACTCAAAGTGTTA |
| Sanger-mut7-R | GAATTAATACTGCAGATGTTG |
| Sanger-DNAH1-mut-F | TTGGCCTTCCAAAGTGCTGG |
| Sanger-DNAH1-mut-R | TTCAGCAGGTTGGCCCTGAC |
| RT-PCR and qPCR: |  |
| *M-Dnah12*-qPCR-F1 | TACAGAGTGTTCTTGTGAAG |
| *M-Dnah12*-qPCR-R1 | TAATTCCTGTTAAGTCCAGC |
| *M-Actb*-qPCR-F1 | CATTGCTGACAGGATGCAGAAGG |
| *M-Actb*-qPCR-R1 | TGCTGGAAGGTGGACAGTGAGG |
| Genotyping: |  |
| *Dnah12-KO-F* | CATTTGGCTTGTTCTTTTGA |
| *Dnah12-KO-R* | CTCTCAAGTGCTGGTATTAC |
| *Dnah12-mut/mut*-F | GCCTAACATTGGCGATGAACT |
| *Dnah12-mut/mut*-R | GCACCAGCACTTATAACTTCAAA |

| **Table S2. The primary antibodies used in this study** | | | | | |
| --- | --- | --- | --- | --- | --- |
| Number | Name | Manufacturer | Cat. Number | Host | Working Conc. |
| 1 | α-Tubulin antibody | Sigma | T6074 | Mouse | 1:200 for IF, 1:3000 for WB |
| 2 | DNAH12 antibody | Origene | TA331519 | Rabbit | 1:100 for IF |
| 3 | DNAH12 antibody | Homemade antibody | Produced in this study | Rat | 1:100 for IF, 1:3000 for WB |
| 4 | DNAH12 antibody | Homemade antibody | Produced in this study | Rabbit | 1:3000 for WB, 2μg for IP |
| 5 | DNAH1 antibody | Abcam | ab122367 | Rabbit | 1:100 for IF |
| 6 | DNAH1 antibody | Homemade antibody | Produced in previous study [4] | Rabbit | 1:3000 for WB |
| 7 | DNALI1 antibody | Sigma | HPA028305 | Rabbit | 1:100 for IF, 1:1000 for WB |
| 8 | DNALI1 antibody | Proteintech | 17601-1-AP | Rabbit | 1:100 for IF, 2μg for IP |
| 9 | SPAG6 antibody | Proteintech | 12462-1-AP | Rabbit | 1:100 for IF, 1:1000 for WB |
| 10 | DNAH17 antibody | Homemade antibody | Produced in previous study  [11] [41] | Rabbit | 1:100 for IF |
| 11 | ODF2 antibody | Proteintech | 12058-1-AP | Rabbit | 1:100 for IF |
| 12 | GAPDH antibody | Proteintech | 60004-1-Ig | Mouse | 1:3000 for WB |
| 13 | ACTB antibody | Abcam | ab8227 | Rabbit | 1:3000 for WB |
| 14 | SPEF2 antibody | Sigma | HPA040343 | Rabbit | 1:100 for IF |
| 15 | DNAI1 antibody | Abcam | ab171964 | Rabbit | 1:1000 for WB |
| 16 | DNAI2 antibody | Proteintech | 17533-1-AP | Rabbit | 1:100 for IF, 1:2000 for WB |
| 17 | Lectin PNA | Thermo Fisher | L32458 | Peanut | 1:200 for IF |
| 18 | mCherry antibody | Abcam | ab167453 | Rabbit | 1:3000 for WB, 2μg for IP |
| 19 | GFP antibody | Abmart | M20004M | Mouse | 1:5000 for WB |
| 20 | Rabbit IgG antibody | Proteintech | 30000-0-AP | Rabbit | 2 ug for IP |
| 21 | RSPH1 antibody | Sigma | HPA017382 | Rabbit | 1:100 for IF, 1:1000 for WB |
| 22 | RSPH9 antibody | Proteintech | 23253-1-AP | Rabbit | 1:1000 for WB |
| 23 | DNAJB13 antibody | Proteintech | 25118-1-AP | Rabbit | 1:1000 for WB |
| 24 | β-Tubulin antibody | Immunoway | YM3030 | Mouse | 1:3000 for WB |

| **Table S3. The secondary antibodies used in this study** | | | | | |
| --- | --- | --- | --- | --- | --- |
| Number | Name | Manufacturer | Cat. Number | Host | Working Conc. |
| 1 | Mouse (Alexa-488) | Molecular Probes | A21121 | Goat | IF: 1:100 |
| 2 | Rabbit (Alexa-555) | Molecular Probes | A31572 | Donkey | IF: 1:200 |
| 3 | Rat (Alexa-488) | Abcam | ab150153 | Donkey | IF: 1:100 |
| 4 | Rat (Alexa-568) | ThermoFisher | A11077 | Goat | IF: 1:200 |
| 5 | HRP Goat anti-Mouse IgG | Biolegend | 405306 | Goat | 1:10000 for WB |
| 6 | HRP Donkey anti-Rabbit IgG | Biolegend | 406401 | Donkey | 1:10000 for WB |
| 7 | HRP Goat anti-Rat IgG | Biolegend | 405405 | Goat | 1:10000 for WB |
| 8 | Clean-Blot™ IP | ThermoFisher | 21230 | - | 1:100 for WB |

**Table S4. Primers used for the construction of plasmids in in-vitro Co-IP assays**

| Primer names | Primer sequences (5'-3') |
| --- | --- |
| p-mCherry / GFP-Gene-vector-F | AGCGGCCGCGACTCTAGATC |
| p-mCherry / GFP-Gene-vector-R | CTTGTACAGCTCGTCCATGC |
| mCherry-m-DNAH12-Stem-F | GCATGGACGAGCTGTACAAGATGTCAGATCCTAACAAAAC |
| mCherry-m-DNAH12-Stem-R | GATCTAGAGTCGCGGCCGCTAAAATCTGTATCATGTGAGACA |
| GFP-m-DNALI1-FL-F | GCATGGACGAGCTGTACAAGATGATACCCCCAGCAGACTC |
| GFP-m-DNALI1-FL-R | GATCTAGAGTCGCGGCCGCTTCACTTCTTCGGTGCGATAA |

| **Table S5. List of candidate dynein protein interactors of DNAH12** | | | | | |
| --- | --- | --- | --- | --- | --- |
| Rank | Gene name | Protein name | seq count  (Anti-DNAH12) | seq count  (Anti-IgG) | Ratio of  DNAH12/IgG |
| 1 | *Dnah12* | DNAH12 | 242 | 0 | - |
| 2 | *Dnali1* | DNALI1 | 30 | 0 | - |
| 3 | *Dnah1* | DNAH1 | 13 | 0 | - |
| 4 | *Dnah3* | DNAH3 | 12 | 0 | - |
| 5 | *Dnah2* | DNAH2 | 7 | 0 | - |
| 6 | *Dnah5* | DNAH5 | 7 | 0 | - |
| 7 | *Dnah17* | DNAH17 | 5 | 0 | - |
| 8 | *Dnah8* | DNAH8 | 18 | 11 | 1.63 |
